# Activation of Piezo1 Inhibits Kidney Cystogenesis

**DOI:** 10.1101/2024.05.11.593717

**Authors:** Qingfeng Fan, Mohamad Hadla, Zack Peterson, Grace Nelson, Hong Ye, Xiaofang Wang, Jean Marc Mardirossian, Peter C. Harris, Seth L. Alper, Y.S. Prakash, Arthur Beyder, Vicente E. Torres, Fouad T. Chebib

## Abstract

The disruption of calcium signaling associated with polycystin deficiency has been proposed as the primary event underlying the increased abnormally patterned epithelial cell growth characteristic of Polycystic Kidney Disease. Calcium can be regulated through mechanotransduction, and the mechanosensitive cation channel Piezo1 has been implicated in sensing of intrarenal pressure and in urinary osmoregulation. However, a possible role for PIEZO1 in kidney cystogenesis remains undefined. We hypothesized that cystogenesis in ADPKD reflects altered mechanotransduction, suggesting activation of mechanosensitive cation channels as a therapeutic strategy for ADPKD. Here, we show that Yoda-1 activation of PIEZO1 increases intracellular Ca^2+^ and reduces forskolin-induced cAMP levels in mIMCD3 cells. Yoda-1 reduced forskolin-induced IMCD cyst surface area *in vitro* and in mouse metanephros *ex vivo* in a dose-dependent manner. Knockout of polycystin-2 dampened the efficacy of PIEZO1 activation in reducing both cAMP levels and cyst surface area in IMCD3 cells. However, collecting duct-specific *Piezo1* knockout neither induced cystogenesis in wild-type mice nor affected cystogenesis in the *Pkd1*^*RC/RC*^ model of ADPKD. Our study suggests that polycystin-2 and PIEZO1 play a role in mechanotransduction during cystogenesis *in vitro*, and *ex vivo*, but that *in vivo* cyst expansion may require inactivation or repression of additional suppressors of cystogenesis and/or growth. Our study provides a preliminary proof of concept for PIEZO1 activation as a possible component of combination chemotherapy to retard or halt cystogenesis and/or cyst growth.

## Introduction

Autosomal dominant polycystic kidney disease (ADPKD) is caused by mutations in *PKD1* or *PKD2*, respectively encoding the proteins polycystin-1 (PC1) and polycystin-2 (PC2). ADPKD is characterized by focal evaginations of tubular epithelium that become disconnected blind sacs. The growth of these blind sacs into fluid-filled cysts slowly disrupts renal architecture and function, ultimately leading to kidney failure in ∼50% of patients.(1) PC1 is a large membrane glycoprotein with 11 transmembrane domains, a large amino-terminal region, and a carboxy-terminal region of 226 amino acids.(2) PC2 is a nonselective cation channel with significant Ca^2+^ permeability.(3) PC1 and PC2 interact physically and function as an asymmetric 1:3 heteromeric complex in the plasma membrane of primary cilia.(4) Reduction or loss of the functional PC complex leads to decreased [Ca^2+^]_i_, resulting in elevated of cyclic AMP (cAMP) levels and a switch of the proliferative response to cAMP from inhibition to stimulation. (1, 5) While several *in vivo* studies support the hypothesis that disruption of [Ca^2+^]_i_ signaling is central to ADPKD pathogenesis, the role of polycystins in regulating [Ca^2+^]_i_ remains elusive. Polycystins have been suggested to play a role in mechanotransduction (6-14), as suggested by their subcellular localizations and physical interactions with various cellular components such as primary cilia,(15,16), cell-cell adherens junctions,(17,18) cell-matrix focal adhesion complexes,(19-23) actomyosin cytoskeleton(13, 24-26), and mechanosensitive cation channels such as TRPC1,(27-29) TRPC3,(30) TRPC7,(30) TRPV4(12,31) and PIEZO1.(6,32) While mechanosensing appears important for cyst generation and growth, the identities of mechanosensors that drive this process remain undefined.

The large mechanosensitive cation channel PIEZO1 is directly gated by membrane tension to mediate Ca^2+^ influx in numerous mammalian cell types.(33-40) PIEZO1 is essential to transduction of distinct mechanical stimuli, such as touch and pain sensation, hearing, and blood pressure regulation.(33, 39, 41-44) In adult mice, PIEZO1 is highly expressed in the lung, bladder and kidney, whereas the homologous mechanosensor PIEZO2 is expressed at lower levels in kidney. (32, 45) PIEZO1 is expressed in the renal tubules, where it can co-immunoprecipitate with PC2.(32, 46) By allowing calcium influx in response to external forces, Piezo1 tightly regulates epithelial cell development, proliferation, and extrusion.(47, 48) Murine PIEZO1 plays roles in vascular smooth muscle arterial remodeling, blood pressure control mediated by flow-induced ATP release from endothelial cells, (49) pressure-sensing in bladder urothelium, (50) and collecting duct osmoregulation.(51)

Peyronnet et al. demonstrated that PC2 regulates stretch activation of PIEZO1, showing requirement of the PC2 N-terminal domain for inhibitory interaction with PIEZO1. (32) PIEZO1 was also critically required for stretch-activated non-selective cationic channel (SAC) activity in collecting duct principal cells and was implicated in urinary osmoregulation.(51) These findings encouraged further investigation of the possible pathophysiological roles of PIEZO1 in ADPKD, a disease associated with increased intrarenal pressures. We hypothesized that cystogenesis in ADPKD results from a failure of mechanotransduction, such that activation of Ca^2+^-permeable mechanosensory channels might be a therapeutic strategy for ADPKD treatment.

In this study, we test the role of Piezo1 in ADPKD cystogenesis. We demonstrate that pharmacological activation of PIEZO1 in mouse inner medullary collecting duct (mIMCD3) cells rapidly increases cytoplasmic [Ca^2+^]_I_, reduces cAMP levels and decreases cystogenesis *in vitro* and *ex vivo*. However, Piezo1 deletion in the collecting ducts neither induced cystogenesis in wild-type mice nor modulated cystogenesis in the *Pkd1*^*RC/RC*^ mouse model of ADPKD, The results suggest redundancy in the mechanosensory pathways of renal tubular epithelial cells, and that PIEZO1 may play context-dependent roles, protective and/or pathogenic.

## Materials and Methods

### Cell culture

Wild-type (WT) and PKD2^-/-^ mouse inner medullary collecting duct (IMCD3) cells (a gift from the Kleene lab)(52) were cultured in DMEM/F12 (1:1) (Gibco/Thermo Fisher Scientific, Waltham, MA) supplemented with 5% fetal calf serum (Sigma-Aldrich, St. Louis, MO) and 1% Pen/Strep (Gibco). WT mouse IMCD cells stably expressing either GCaMP6s (53) or jRgeco1a(54) together with flamindo2-GFP(55) were cultured in the same medium supplemented with G418 (500 μg/ml).

### Cyclic AMP ELISA assay

cAMP was measured by ELISA (Direct cAMP ELISA kit) per manufacturer’s instructions (Enzo Life Sciences, Farmingdale, NY). WT and PKD2^-/-^ mouse IMCD cells cultured in 6-well plates and treated as indicated were homogenized in 300 μl of 0.1 M HCl per well at room temperature. Protein concentrations were measured (Pierce BCA Protein Assay Kit, Thermo Fisher Scientific). After lysate centrifugation at 12,000 rpm for 15 minutes at 4°C, supernatant optical density at 405 nm was measured by SYNERGY HTX Multi-Mode Microplate Reader (BioTek, Winooski, VT), with 4-6 wells per condition in each of three independent experiments. Results are presented as pmol/mg of protein. Excess supernatants were stored at -80°C for later analysis.

### Live cell imaging

GCaMP6s (Addgene plasmid #40753(53)) binds Ca^2+^ with peak emission at 510 nm when excited at 480 nm.(53) WT mouse IMCD cells stably expressing GCaMP6s were cultured in a 35mm dish with a glass bottom. Live images were taken at 3 seconds intervals using a Zeiss confocal microscope equipped with a 40x oil objective (Carl Zeiss Vision Inc., San Diego, CA). After baseline imaging, PIEZO1 agonist Yoda-1 was added to final concentrations of 250 or 750 nM. Of note, the EC_50_ of Yoda1 for [Ca^2+^]_I_ elevation in HEK293T cells respectively expressing murine or human PIEZO1 was reported as 17.1 and 26.6 μM.(56) GCaMP6 fluorescence intensity (FI) of the region-of-interest (ROI) relative to baseline (F/F0) was calculated through the Zeiss software. WT mouse IMCD3 cells stably expressing [Ca^2+^] indicator jRgeco1a and cAMP inverse indicator flamindo2-GFP were cultured in a 35mm dish with a glass bottom.(55) After baseline imaging, cells were treated with 750 nM Yoda-1, 10 μM forskolin, or forskolin plus Yoda-1 as indicated. GCaMP FI relative to baseline (F/F0) was automatically calculated to estimate relative change in [Ca^2+^]_i_, whereas flamindo2-GFP F0/F reflected the change in cAMP levels.

### 3D culture and *in vitro* cystogenesis assay

WT and Pkd2^-/-^ mouse IMCD cells were cultured in Matrigel for *in vitro* cystogenesis analyses. 40 μl per well of 1:1 cold matrigel/growth media solution (1:1) was added to a pre-chilled 96-well plate, and Matrigel was allowed to polymerize at 37°C for 1 hour. 5000 cells per well in 150 μl growth media were seeded and incubated at 37°C for 90 min. Immediately after gentle aspiration of 130 μl media, 40 μl of 50% Matrigel was added and cells were incubated 1 h at 37°C. After further addition of 150 μl growth media was added and cells were restored to incubators. Compounds or vehicle were added daily with medium changes. Plates were imaged within a 4-hour interval in a multi-spheroid mode using an Incucyte S3 high-content screening system (Sartorius, Bohemia, NY). Average cyst areas (μm^2^) were automatically measured and analyzed with the built-in Live-Cell Analysis System (Sartorius). Samples were processed in 8-16 wells per condition in each of three independent experiments.

### Animal models

All experimental protocols were approved by the Institutional Animal Care and Utilization Committee of Mayo Clinic (Rochester, MN). *Pkd1*^RC/RC^ and *Pkhd1*-Cre (expressing Cre recombinase specifically in collecting duct) mouse models used in this study were inbred on a C57Bl/6 background. B6.Cg-*Piezo1*^tm2.1Apat^/J (designated here as *Piezo1*^fl/fl^) mice, inbred on a C57Bl/6 background, were from JAX (stock # 029213, originally donated by the Patapoutian laboratory).(57) *Piezo1*^fl/fl^, *Pkd1*^RC/RC^, and *Pkhd1*-Cre mice were crossed to generate *Pkhd1*-Cre; *Piezo1*^fl/fl^ and *Piezo1*^fl/fl^; *Pkd1*^RC/RC^; *Pkhd1*-Cre mice. All mice were euthanized at 4 months of age. Tissue samples for genotyping were collected by tail-clipping at 2 weeks of age. Genomic DNA was extracted using Fast DNA Extraction Method.(58) *Piezo1, Pkhd1*-Cre and *Pkd1* genotyping was performed by PCR using appropriate primers: *Pkhd1*-Cre forward-aggttcgttcactcatgga and reverse-tcgaccagtttagttaccc; Piezo1 forward-gcctagattcacctggcttc and reverse-gctcttaaccattgagccatct; *Pkd1*^RC/RC^ forward-caaaggtctgggtgataactggtg and reverse-caggacagccaaatagacaggg.

Twenty-four-hour urine output was collected in metabolic cages one day before sacrifice at 4 months of age (ketamine 90 mg/kg and xylazine 10 mg/kg IP). Blood was obtained by cardiac puncture. The left kidney and part of the liver were placed into pre-weighed vials containing 10% formaldehyde in phosphate buffer and subsequently weighed. The right kidney and remaining liver were immediately frozen in liquid nitrogen and weighed.

### Histology and histomorphometric analysis

5 μ paraffin-embedded kidney tissue sections were stained with hematoxylin-eosin. Images were taken using a Nikon SMZ800 microscope equipped with a color digital DS-Ri1 camera and NIS-Elements BR Imaging Software version 4.30 (Nikon, Minato City, Tokyo, Japan). Cyst area was analyzed with Meta-Morph software (Universal Imaging, West Chester, PA). A standardized threshold separated cysts from non-cystic tissue to calculate the renal cystic index (%), the ratio of cyst area to total kidney area.

### Immunohistochemistry staining

Immunohistochemistry was performed on 5 μm paraffin-embedded kidney tissue sections. Deparaffinized and rehydrated tissue sections were incubated in 1.5% H_2_O_2_ for 20 minutes to block endogenous peroxidase, followed by antigen retrieval in 1 mM EDTA (pH 8.0) for 45 minutes in a vegetable steamer. Block was performed in 10% goat serum for 1 hour at room temperature to reduce non-specific background. Rabbit anti-AQP2 (1:200; cat. No. NB110-74682, Novus Biologicals, Centennial, CO) and rabbit anti-Piezo1 (1:200; cat. no. 15939-1-AP, Proteintech Group, Inc., Rosemont, IL) antibodies were incubated with tissue sections overnight at 4°C. After 3 washes with 1x DPBS, HRP-conjugated goat anti-rabbit IgG antibody (ab214880, Abcam, Waltham, MA) was applied for 45 minutes at room temperature. After 3 washes, a DAB substrate solution (Sigma-Aldrich) was added, and color was developed and monitored under the microscope, followed by counterstaining with hematoxylin. Rapid dehydration was performed in 100% ethanol, followed by 3 changes of Histo-clear solution. Slides were mounted with Permount. Digital images were acquired using a light microscope with a high-resolution Nikon digital camera DXM1200 (Nikon) and Meta-Morph software (Universal Imaging).

### Metanephric culture and *ex vivo* cystogenesis assay

Metanephric kidneys were dissected at embryonic day E13.5, placed on a 0.4 μm transparent membrane (Greiner Bio-One, Monroe, NC), and cultured in DMEM/F12 (1:1) supplemented with 2.8 nM selenium, 3 ng/ml triiodothyronine, 5 μg/ml insulin, 5 μg/ml transferrin, 25 ng/ml prostaglandin E1 and 1% Pen/Strep at 37°C in 5% CO_2_. Paired kidneys were treated daily with vehicle, Yoda-1 (750 nM), forskolin (10 μM), or forskolin (10 μM) plus Yoda-1 (750nM), with daily replacement of media. Metanephroi were imaged daily by Nikon SMZ800 microscope equipped with a color digital DS-Ri1 camera and NIS-Elements BR Imaging Software version 4.30 (Nikon). Images were analyzed using an automated Metanephric Analysis Toolkit v0.3 that segments kidneys, detects the cysts, and measures total and cystic areas. The cystic index was calculated (% cyst area relative to intact metanephros area).

### Statistical methods

Each experimental group included 9–10 animals. Ten male and ten female animals in each experimental or treatment group provide 90% power to detect a reduction in cystic index from 25% to 20% assuming 25±5% control group cyst volume. Data are presented as means ± SD. Comparisons were performed by one-way or two-way ANOVA with Graphpad Prism 9 (Graphpad Software, LLC., San Diego, CA). A p-value <0.05 was considered to represent statistical significance.

## Results

### PIEZO1 agonist Yoda1 inhibits forskolin-induced increases in. cAMP levels

WT mouse IMCD cells respond to adenylyl cyclase agonist forskolin (59) (10 μM) with rapidly increased cAMP levels persisting for 3 h (**Figure 1A**). Forskolin-induced cAMP increase was less pronounced at 15 and 30 minutes in PKD2^-/-^ than in WT IMCD cells (**Figure 1C**). Whereas 750 nM Piezo1 agonist Yoda1 (56) did not influence cAMP levels, Yoda1 pretreatment for 1 h attenuated the forskolin-induced increase in cAMP In both WT and PKD2^-/-^ IMCD cells (**Figure 1B,C**). The absence of PC2 also dampened the efficacy of Yoda1 in reducing cAMP levels, (**Figure 1C,D**) suggesting possible modulation of PIEZO1 function by PC2 expression.

**Fig 1.**
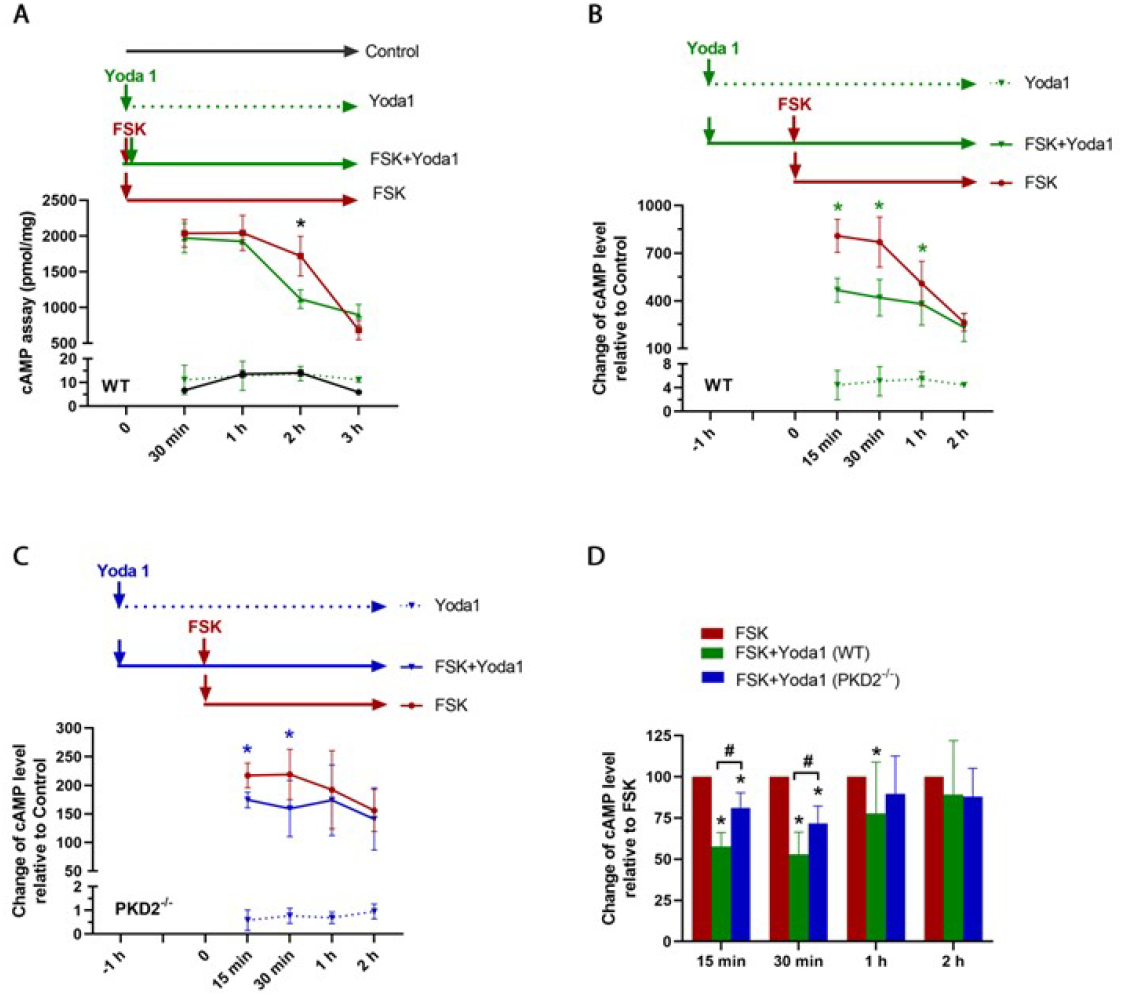
Activation of Piezo1 through Yoda1 inhibits increase of cAMP induced by forskolin. **A**. cAMP was measured in Wild type (WT) mouse inner medullary collecting duct (IMCD) cells treated with 10 μM forskolin (FSK), 750 nM Yoda1, or FSK plus Yoda1. *Two-way ANOVA*, ^*\**^*p<0*.*05 FSK+Yoda1 vs FSK alone*. **B & C**. cAMP was measured in WT **(B)** and PKD2^-/-^ **(C)** mouse IMCD cells pretreated 1 h with 750 nM Yoda1 then exposed for the indicated times to 10 μM FSK. Untreated cells served as controls. Change in cAMP level relative to control is presented. *Two-way ANOVA*, ^*\**^*p<0*.*05 FSK+Yoda1 vs FSK alone*. **D**. Change in cAMP level relative to FSK treatment is presented for data shown in B and C. *One-way ANOVA*, ^*\**^*p<0*.*05 FSK+Yoda1 vs FSK alone; Two-way ANOVA*, ^*#*^*p< 0*.*05 PKD2*^*-/ -*^ *vs WT*.

### Activation of PIEZO1 increases cytosolic calcium levels and dampens the increase of cytosolic cAMP induced by forskolin

Dynamics of intracellular calcium and cAMP were monitored by confocal live cell imaging of IMCD cells stably transfected with Ca^2+^ indicator GcaMP6s or cotransfected with dual Ca^2+^/cAMP indicators jRgeco1a and reverse Flamindo2-GFP. Yoda 1 addition rapidly increased intracellular free calcium [Ca^2+^]_i_ in a dose-dependent manner (**Figure 2A**). In IMCD cells stably expressing both [Ca^2+^]_i_ indicator jRgeco1a and cAMP indicator reverse Flamindo2-GFP, Yoda1’ stimulation-induced [Ca^2+^]_i_ increase was accompanied by immediate but short-lived cAMP decrease (**Figure 2B**). Forskolin induced an immediate decrease of cytosolic cAMP which returned to baseline within ∼4 min. Yoda1 pretreatment greatly enhanced the forskolin-induced decrease in cAMP to slowly oscillate over the next 60 min (**Figure 2C**). Yoda1 pretreatment substantially attenuated and stabilized the forskolin-induced stable decline in cAMP levels. (**Figure 2D**).

**Fig 2.**
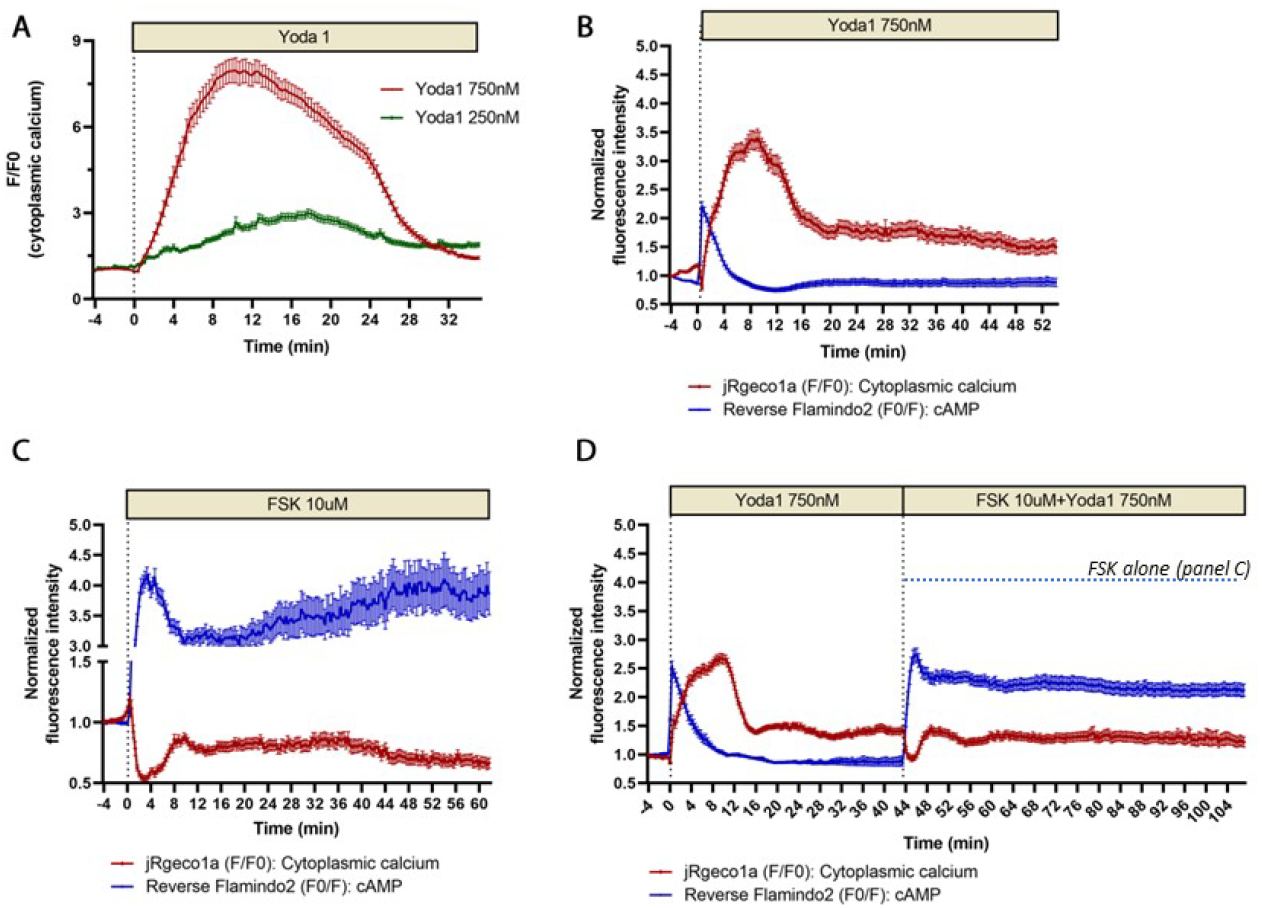
Activation of Piezo1 increases cytosolic Ca^2+^ levels and inhibits increase of cytosolic cAMP induced by forskolin. **A**. Wild type (WT) mouse inner medullary collecting duct (IMCD) cells stably expressing GCaMP6s, a GFP-based fluorescent Ca^2+^ indicator were treated with 250 nM or 750 nM Yoda1. Relative Fluorescence intensity (F) relative to baseline (F0) was proportional to [Ca^2+^]i. **B-D**. WT mouse IMCD cells stably expressing red fluorescent Ca^2+^ sensor protein jRgeco1a and green fluorescent cAMP sensor Flamindo2-GFPwere treated with 750 nM Yoda1 alone (B), 10 μM forskolin (FSK) alone **(C)**, or 750 nM Yoda1 followed by 10 μM FSK **(D)**. Fluorescence intensity (F) normalized to the baseline (F0) is presented. Note that Flamindo2 fluorescence emission is inversely proportional to cAMP levels.

### Activation of Piezo1 decreases in vitro cellular cystogenesis induced by forskolin

Forskolin is known to promote cystogenesis through elevation of cAMP levels.(5) We found that forskolin treatment of IMCD cells cultured in Matrigel gradually induced cystogenesis, yielding mean day 6 WT cyst area of 11,000 μm^2^ and mean PKD2^-/-^ cyst area 5,000 μm^2^. In WT cells, Yoda1 treatment of WT cells inhibited forskolin-induced cyst growth in a dose-dependent manner from ≈18% (250 nM Yoda1) to 84% (for 1000 nM Yoda1) **Figure 3A,B**). The forskolin-induced 280% increase cyst area in WT IMCD cells (day 6) was nearly abrogated in PKD2^-/-^ IMCD cells (**Figure 3C,B**), whereas combined exposure to forskolin and ≥ 500 nM Yoda1 reduced both WT and PKD2^-/-^ cyst surface area to control values or lower (**Figure 3C**). The dose-dependent Yoda1-induced reduction in forskolin-stimulated cyst area in WT cells (∼45%) was less prominent in PKD2^-/-^ (30%, p< 0.001) (**Figure 3D**), suggesting that PIEZO1 activation-associated reduction in forskolin-stimulated cystogenesis was more effective in the presence than in the absence of PC2.

**Fig 3.**
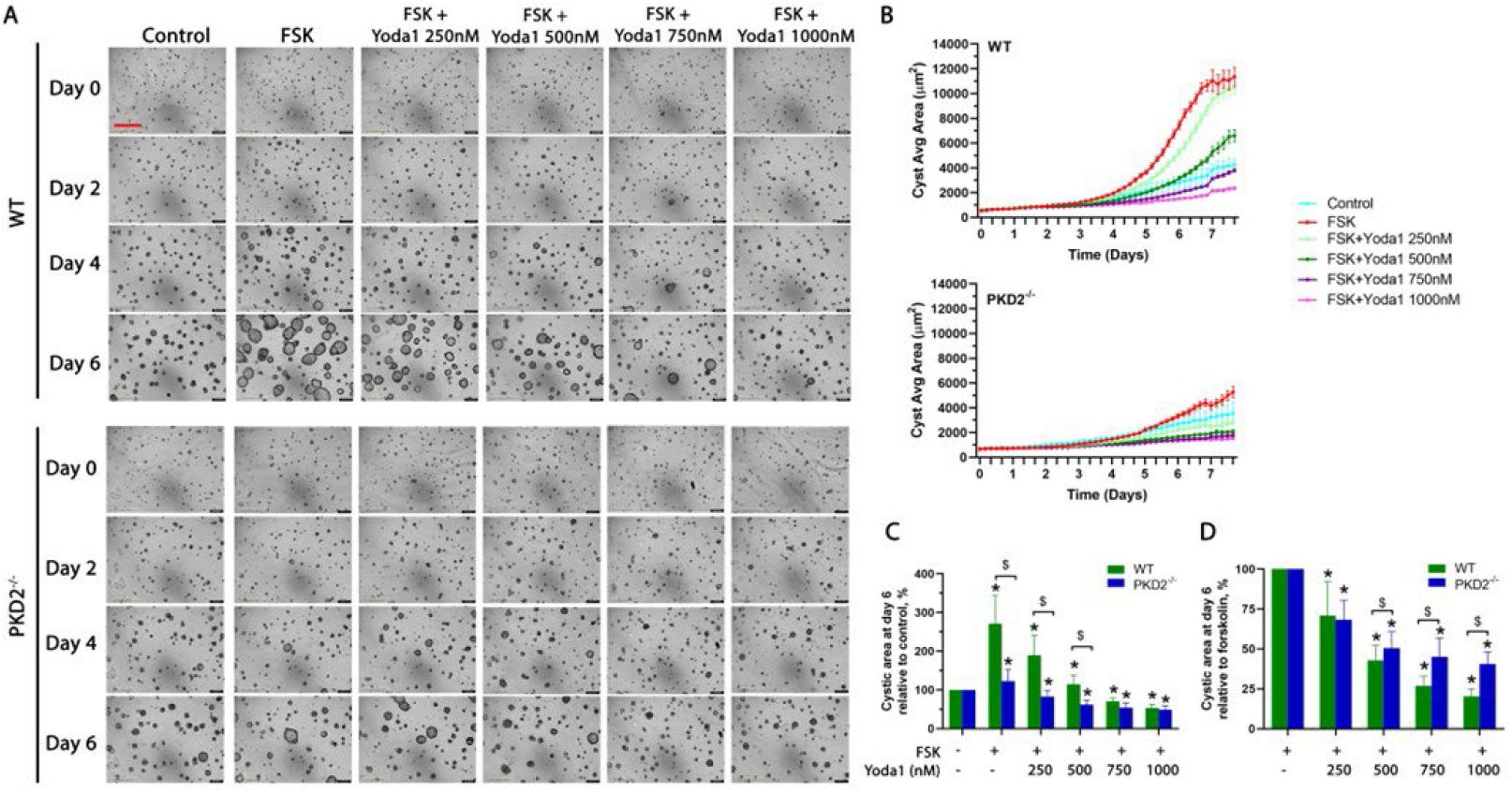
Activation of Piezo1 decreases cellular cystogenesis induced by forskolin. **A**. Wild type (WT) and PKD2^-/-^ mouse inner medullary collecting duct (IMCD) cells cultured in Matrigel were treated with 10 μM forskolin (FSK), or FSK plus Yoda1 during a 3D cystogenesis protocol. *Representative images from 3 independent experiments are presented. Scale (red bar) = 400 μm*. **B**. Mean cyst areas (μm^2^) were measured over 8 days. **C**. Cyst area at day 6 relative to control was compared. *One-way ANOVA*, ^*\**^*p<0*.*0001 FSK or FSK+Yoda1 vs control; Two-way ANOVA*, ^*$*^*p<0*.*0001 PKD2*^*-/-*^ *vs WT*. **D**. Cyst area at day 6 relative to FSK was compared. *One-way ANOVA*, ^*\**^*p<0*.*0001 FSK+Yoda1 vs FSK; Two-way ANOVA*, ^*$*^*p<0*.*0001 PKD2*^*-/-*^ *vs WT*.

### Piezo1 activation decreases forskolin-induced ex vivo cystogenesis in Pkd1^RC/RC^ metanephric organ culture

Cultured embryonic kidneys from *Pkd1*^RC/RC^ mice form renal cysts when chronically stimulated by forskolin. However, the mean cyst area of 53% on day 6 was reduced to 12% in the combined presence of forskolin and Yoda1 (**Figure 4A,B**), representing a 76% inhibition of forskolin-induced cystogenesis (p<0.001) (**Figure 4C**).

**Fig 4.**
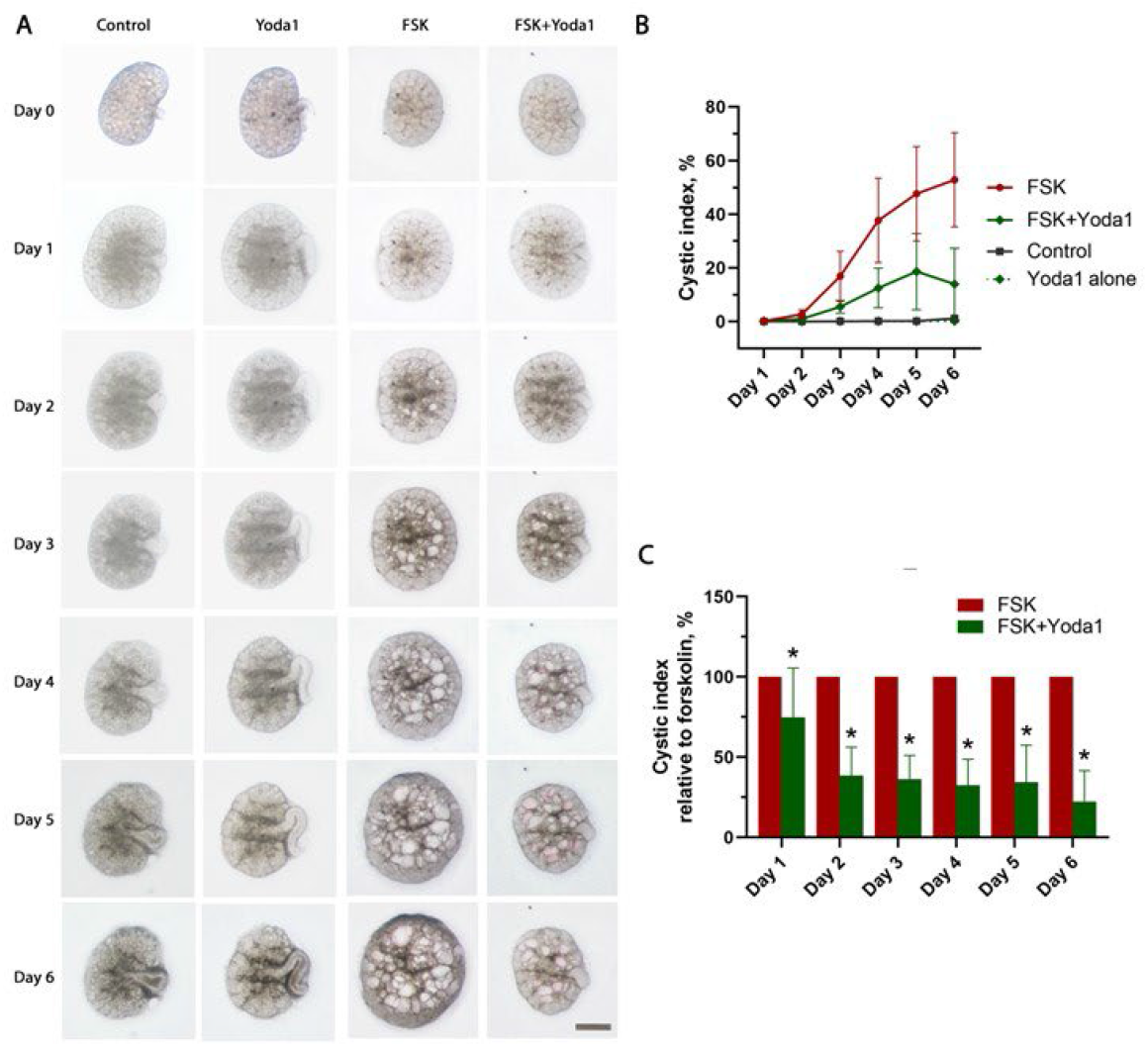
Activation of Piezo1 decreases *ex vivo* cystogenesis induced by forskolin in *Pkd1*^RC/RC^ metanephric organ culture. **A**. E13.5 metanephric kidneys from *Pkd1*^RC/RC^ mice were cultured on 0.4 μm transparent membranes in 6-well plates, and treated with 750 nM Yoda1 alone, 10 μM forskolin (FSK) alone, or 10 μM FSK plus 750 nM Yoda1. *Representative images are presented. Scale = 1000 μm*. **B**. The cystic index, (cyst area relative to intact metanephros area (%), was calculated by automated Metanephric Analysis Toolkit. *n=10 in FSK and FSK+Yoda1; n=5 in Control and Yoda1 alone*. **C**. The cystic index relative to FSK was compared. *One-way ANOVA*, ^*\**^*p<0*.*001 FSK+Yoda1 vs FSK*.

### Piezo1 knockout does not induce renal cystogenesis in WT mice and does not exacerbate renal cystogenesis in Pkd1^RC/RC^ mice

To determine if Piezo1 protects against cystogenesis upstream of polycystins, we generated a renal collecting duct-specific *Piezo1* knockout mouse model (*Piezo1*^fl/fl^; *Pkhd1*-Cre). As global *Piezo1* knockout is embryonic lethal due to defective vasculogenesis,(60) we crossed floxed *Piezo1* mice with *Pkhd1*-cre mice to restrict knockout to collecting duct epithelial cells (expressed in the collecting ducts). We found that neither the ratio of kidney weight to body weight nor the kidney cystic index differed between *Piezo1*^fl/fl^; *Pkhd1*-Cre and WT mice (**Figure 5**). There were no sex differences in mice of either genotype (**Supplemental figure 1**). This indicates that collecting duct knockout of *Piezo1* does not induce cystogenesis.

**Fig 5.**
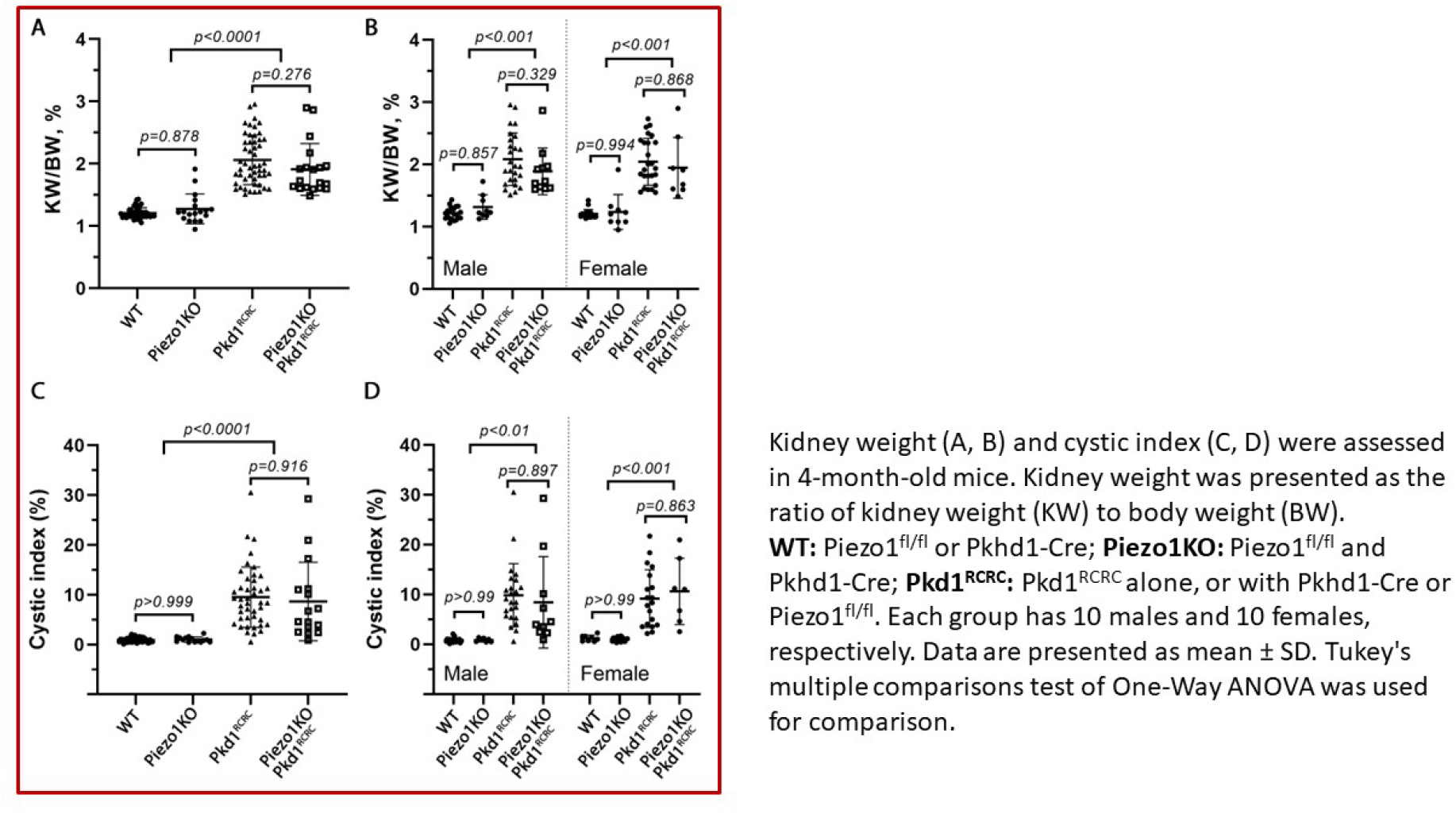
Piezo1 knockout shows no effect on kidney cystogenesis in *Pkd1*^RC/RC^ mice. **A & B**. Ratio of kidney weight (KW) to body weight (BW) in 4-month-old mice of indicated genotypes. **C & D**. Cystic index (%) of the same mice. **WT:** *Piezo1*^fl/fl^ or *Pkhd1*-Cre. **Piezo1KO:** *Piezo1*^*fl/fl*^*;Pkhd1-Cre*. ***Pkd1***^**RC/RC**^: *Pkd1*^RC/RC^ alone, or with *Pkhd1*-Cre or *Piezo1*^fl/fl^. Each group had 10 males and 10 females, respectively. *Data are presented as mean ± SD. Tukey’s multiple comparisons test of One-Way ANOVA was used for comparison. The p values are shown in the graph*.

To determine if *Piezo1* modulates the protective function of polycystins (presumably acting downstream of polycystins), we generated a *Piezo1* knockout mouse in the background of *Pkd1* mutant mice (*Piezo1*^fl/fl^; *Pkhd1*-cre; *Pkd1*^RC/RC^).Collecting duct-restricted *Piezo1* knockout in *Pkd1*^*RC/RC*^ mice affected neither the ratio of kidney weight to body weight nor or the cystic index as compared to *Pkd1*^*RC/RC*^ mice. Thus, knocking out *Piezo1* on an orthologous *Pkd1* mutant background did not exacerbate cystogenesis (**Figure 5**). Again, no sex differences were observed (**Supplemental figure 1**). Collecting duct knockout of *Piezo1* did not affect body weight or urine volume and did not lower the kidney weight to body weight ratio in either WT or *Pkd1*^RC/RC^ mice (**Supplemental figure 2**).

We validated *Piezo1* knockout in the floxed mouse model by immunohistochemistry. Anti-PIEZO1 signal was evident in renal tubules of Piezo1^fl/fl^ (without Cre) (**Supplemental figure 3**), particularly so in regions of AQP2-positivity, especially in cyst epithelial cells (**Figure 6**). However, Anti-PIEZO1 signal intensity was significantly reduced in *Piezo1*^fl/fl^;*Pkhd1*-Cre mice (**Supplemental figure 4**), with total absence from AQP2-positive regions (**Figure 6**). These results strongly suggest successful knockout of *Piezo1* in collecting duct.

**Fig 6.**
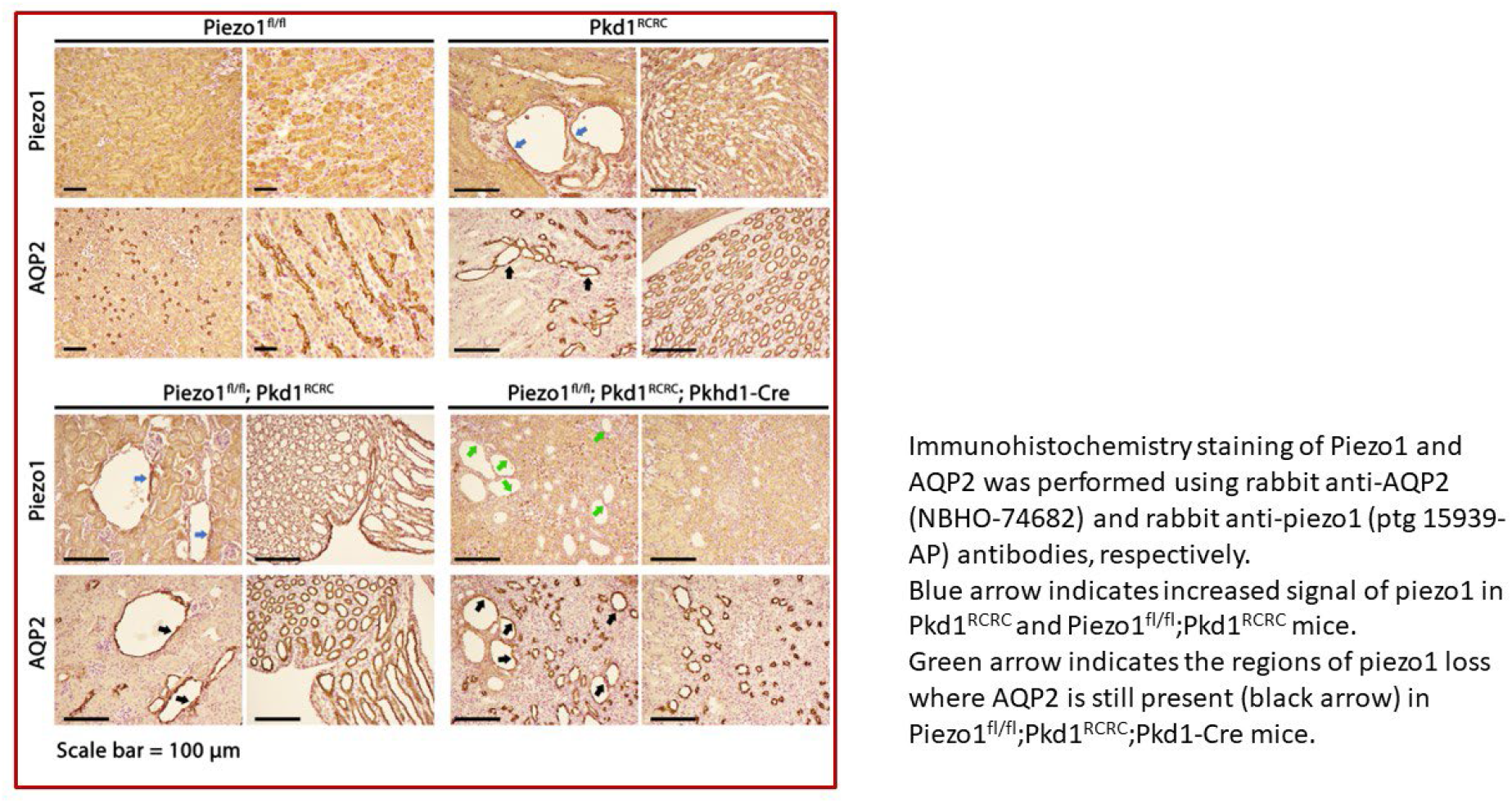
Immunohistochemical localization of kidney PIEZO1 and AQP2. FFPE kidney sections were subjected to immunohistochemical staining of PIEZO1 and AQP2. Blue arrows indicate increased PIEZO1 signal in cystic areas in *Pkd1*^RC/RC^ mice. Sequential sections were stained for *Piezo1*^*fl/fl*^*;Pkd1*^*RC/RC*^ and *Piezo1*^*fl/fl*^*;Pkd1*^*RC/RC*^*;Pkd1-Cre* mice. Blue arrows indicate increased PIEZO1 signal in AQP2-positive regions (black arrows) in *Piezo1*^*fl/fl*^*;Pkd1*^*RC/RC*^ mice. Green arrows indicate regions of PIEZO1 absence in AQP2+ cells (black arrows) in *Piezo1*^*fl/fl*^*;Pkd1*^*RC/RC*^*;Pkd1-Cre* mice. *Representative images from 3-4 mice are presented for each condition. Scale = 100 μm*.

We also performed metanephric organ cultures procured from *Pkd1*^RC/RC^ mice with *Piezo1* knockout. *Piezo1* knockout in embryonic kidneys was strongly supported by Cre PCR (**Supplemental figure 5**). We found that forskolin-induced *ex vivo* cystogenesis was inhibited significantly by Yoda1 in *Pkd1*^RC/RC^ mice, with day 6 cystic index reduced from 48% to 20% (p<0.001) (**Figure 7**). Surprisingly, Yoda1 was also effective in reducing cystic index of *Pkd1*^RC/RC^ metanephric kidneys with *Piezo1* knockout. This could be explained by activation of the Piezo1 channel in tubular cells other than the collecting duct, or by off-target effects of Yoda1. Knockout of collecting duct *Piezo1* failed to alter kidney cystogenesis *Pkd1*^RC/RC^ embryos, as demonstrated by the similar cystic indices in embryonic kidneys of *Piezo1*^fl/fl^;*Pkd1*^RCRC^ and *Piezo1*^fl/fl^;*Pkd1*^RCRC^;*Pkhd1*-Cre mice (**Supplemental figure 6**).

**Fig 7.**
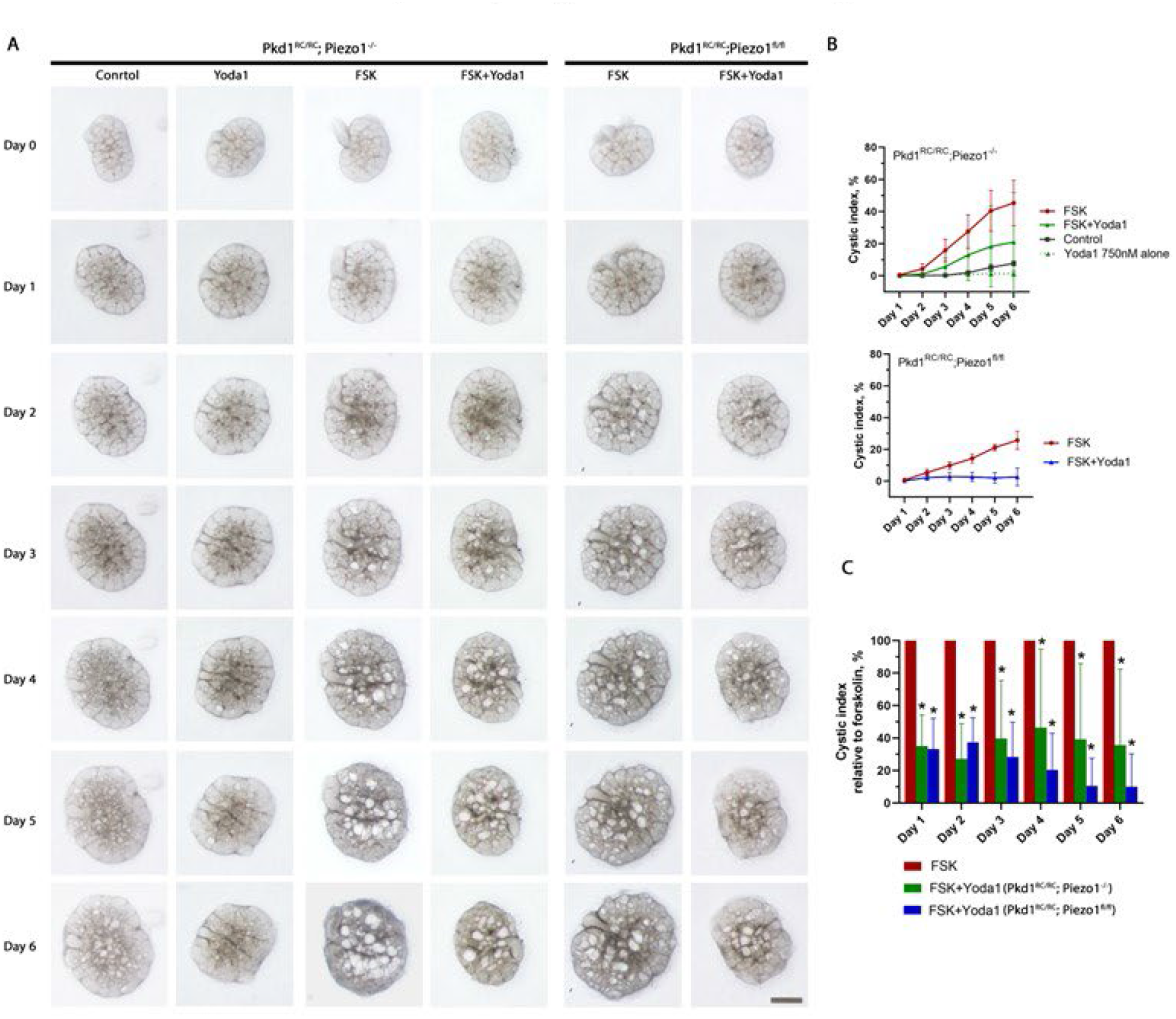
Activation of Piezo1 decreases forskolin-induced ex vivo cystogenesis in *Pkd1*^RC/RC^ mice with and without Piezo1 knockout. **A**. E13.5 metanephric kidneys from Pkd1^RC/RC^ mice with and without *Piezo1* knockout, cultured on 0.4 μm transparent membranes in 6 well plates, were treated with 750 nM Yoda1 alone, 10 μM forskolin (FSK) alone, or 10 μM FSK plus 750 nM Yoda1. *Representative images are presented. Scale = 1000 μm*. **B**. Cystic index of metanephroi of indicated culture age, treatment, and genotype. *n=7 in the groups of FSK and FSK+Yoda1 and n=3 in the groups of Control and Yoda1 alone in Pkd1*^*RC/RC*^*;Piezo1*^*-/-*^ *mice. n=5 in the groups of FSK and FSK+Yoda1 in Pkd1*^*RC/RC*^*;Piezo1*^*fl/fl*^ *mice*. **C**. Cystic index of indicated conditions and genotypes relative to FSK alone. *One-way ANOVA*, ^*\**^*p<0*.*001 FSK+Yoda1 vs FSK*.

## Discussion

In this study we have evaluated the role of PIEZO1 in renal cystogenesis. We found that pharmacological activation of mechanosensitive ion channel PIEZO1 increases [Ca^2+^]_i_, reduces cAMP levels, and inhibits cystogenesis in both *in vitro* and *ex vivo*. These findings are concordant with the phenotypic switch phenomenon in ADPKD, in which elevated [Ca^2+^]_i_ reverses the cystogenic effect of cAMP.(61, 62) However, *Piezo1* knockout in the collecting duct was without measurable effect on induction or modulation of *in vivo* renal cystogenesis.

ADPKD is caused predominantly by mutations in either PC1 or PC2. ADPKD is a devastating condition leading to relentless growth of renal cysts eventuating in renal failure.(63-69) Although the molecular mechanisms responsible for cyst formation have been extensively studied, the function of PC1 remains elusive. Altered [Ca^2+^]_I_ signaling has a central role in ADPKD pathogenesis. (3, 70, 71) ADPKD is one of the most common ciliopathies.(72) The primary cilium and plasma membrane have been described as cellular components devoted to mechanosensing. PC1 and PC2 have been proposed as orchestrators that control cellular mechanosensitivity and regulate [Ca^2+^]_I_ homeostasis. Recent cryoelectron microscopy studies suggest that PC1 and PC2 form the polycystin complex in a 1:3 stoichiometry.(73) The large extracellular domain of PC1 is proposed to have a mechanosensor role in detecting urinary flow through renal tubules. Several studies have suggested that mechanosensation of shear stress at the primary cilium of kidney epithelial cells increases ciliary calcium levels mediated by the polycystin complex.(9, 10, 74-76) However, other studies have challenged the idea of primary cilia as Ca^2+^-responsive and Ca^2+^-regulated mechanosensors, proposing that mechanosensation by the primary cilium might be independent from ciliary calcium.(77) Their conclusion that shear stress-induced Ca^2+^ influx into the cell body diffuses into cilium raises the possibility that PC1 modulates mechanosensation and Ca^2+^ signaling by interacting with other mechanosensitive ion channels. PIEZO1 opens in response to direct physical deformation of the lipid bilayer, such as an increase in lateral membrane tension, and thus obeys the so-called force-from-lipid paradigm established for several mechanosensitive ion channels.(78, 79) Furthermore, renal SACs are strongly dependent on PIEZO1 expression and PC2 negatively regulates the stretch sensitivity of these types of channels.(32)

Our study supports a role for PIEZO1 activation in alleviating cystogenesis. Pharmacological activation of PIEZO1 reduced cAMP levels and concomitantly increased [Ca^2+^]. PIEZO1 activation also inhibited cyst formation > 75% in both *in vitro* and *ex vivo* assays. The effect of PIEZO1 activator Yoda1 is consistent with reversing the phenotypic switch, such that reduction in [Ca^2+^]_i_ associated with expression of mutant PC1 or PC2 switches the cellular response to cAMP from inhibition to stimulation of proliferation. Our study further confirms that a molecule that both increases [Ca^2+^]_i_ and reduces cAMP has a favorable anti-cystogenic profile.

Knockout of *Piezo1* in renal collecting ducts, however, failed to induce renal cystogenesis in WT mice and did not exacerbate renal cystogenesis in a *Pkd1* orthologous mouse model. One explanation is possible redundancy in mechanotransduction signaling (80, 81), such that *Piezo1* knockout was compensated by other collecting duct mechanosensors such as TRPV4. In fact, the role of PIEZO1 in in vitro and ex vivo cystogenesis has many resemblances to previously described roles of TRPV4. TRPV4 forms a mechanosensitive heterometric complexes with PC2, (12) and TRPV4 activator diminishes cyst development and growth in PCK rats.(82) However, *TRPV4* knockdown does not result in a cystic phenotype in mice or zebra fish. (12) Interestingly, PIEZO1 can act upstream of TRPV4 to induce pathological changes in endothelial cells due to shear stress.(83) PIEZO1 activation by fluid shear stress initiates a Ca^2+^_i_ signal that activates TRPV4, which in turn mediates the sustained phase Ca^2+^ elevation that triggers endothelial cell pathology. Thus, deleterious effects of shear stress are initiated by Piezo1 but require TRPV4.(83) Alternatively, a PIEZO1 role in cystogenesis might impact renal tubular segments other than collecting duct. While most renal cysts appear to originate from the collecting ducts,(84) cysts may also derive from other nephron segments, particularly in embryonic state (85), perhaps explaining the anti-cystogenic effect of Yoda1 in metanephric organ culture from *Pkd1*^RC/RC^ mice, despite PIEZO1 knockout in collecting ducts. Alternatively, Yoda1’s anti-cystogenic effect in metanephroi might represent one or more off-target effects of the drug.

Overexpression of PC2 impaired native SACs in renal tubular epithelial cells.(32) The inhibitory effect of PC2 on native SACs could be reversed by overexpressing PC1, while it was mimicked by *Pkd1* deletion. (32) PC2 co-immunoprecipitated with Piezo1 and deletion of its N-terminal domain prevented both this interaction and inhibition of SAC activity. These findings suggest that renal SACs depend on or are regulated by PIEZO1 but are critically conditioned by the PC1/PC2 ratio. (13, 86) This important ratio between PC1 and PC2 would potentially explain our finding that the absence of PC2 in IMCD cells dampened both the efficacy of Yoda1 in reducing cAMP levels and in reducing cyst area.

PIEZO1 plays an important role during early development, and its constitutive genetic deletion is embryonic lethal.(60, 87) Notably, specific Piezo1 knock-out in the endothelium profoundly alters its sensitivity to shear stress, as well as the vascular architecture.(87) Martins et al. have reported conditional knockout of *Piezo1* targeted to renal epithelial cells of the collecting ducts and renal tubules. Induction began at 8 weeks of age and continued for 6 weeks before mice were studied. (51) PIEZO1 transcript expression in inner medulla was significantly but only partially reduced. The authors did not report kidney weights or anatomy. However, a role for PIEZO1 in regulation of urine osmolarity post-dehydration or fasting was reported. The mechanism remains unknown.(51)

Interestingly, immunohistochemical staining revealed remarkable increases in PIEZO1 abundance in *Pkd1*^RC/RC^ mouse kidney, indicative of a possible compensatory role of Piezo1. Nuclear translocation of Piezo1 has been reported in cells from dilated tubules. In fact, mutant *Camsap3*^dc/dc^ transgenic mice expressing a truncated CAMSAP3 protein with a deleted C-terminal microtubule-binding domain in proximal renal tubules developed significant cystic disease.(88) This member of the calmodulin-regulated spectrin-associated protein (CAMSAP)/Nezha/Patronin family is essential for epithelial microtubule organization. In *Camsap3* mutant mice, nuclear PIEZO1 was significantly increased both in cells from dilated and non-dilated tubules, but nuclear YAP was increased only in cells of dilated tubules. The transcriptional activator YAP is a downstream effector of the Hippo pathway that regulates cell proliferation as well as mechanical tension–dependent cell shape regulation.(89, 90) In *Pkd1* mutant mice, the transcriptional co-activator YAP/TAZ and its effector c-Myc are activated, thereby promoting cystogenesis.(91) Additional reports have shown that the relocalization of PIEZO1 from cytoplasm to nuclear region in non-dilated cells suggests that this mechanosensor might begin to be activated prior to the detectable deformation of cells, most likely in response to cell ‘stretch’.(48) Nuclear translocation of both YAP and Piezo1 in stretched cells suggests that Piezo1 may cooperate with YAP in promoting cyst formation and enhancing proliferation in response to cell stretch or other mechanical forces.

Our study reports concomitant measurement of live cell [Ca^2+^]_i_ and intracellular cAMP using stably transfected, genetically encoded biosensors. Optimization of this method would accelerate high throughput screening of molecules that could increase intracellular calcium and reduce cAMP for ADPKD treatment. *in vitro* 3D cystogenesis assays and metanephric organ cultures have been used as systems for evaluating therapeutic agents.(92, 93) These assays, however, share the drawback of a requiring forskolin exposure for cyst formation. Another limitation to our study is the use of pharmacological PIEZO1 activation which does not recapitulate physiological conditions, but does provide proof of concept. In vivo use of Yoda1 was not attempted due to concerns about off-target activity and toxicity.

In conclusion, we demonstrate that PIEZO1 activation inhibits cystogenesis both *in vitro* and *ex vivo*, likely by increasing [Ca^2+^]_i_ and decreasing cAMP levels. Absence of PC2 dampened efficacy of Yoda1 in reducing both cAMP levels and cyst area. PIEZO1 deletion in collecting duct did not induce cystogenesis in wild-type mice and did not exacerbate cystogenesis in an orthologous *Pkd1* murine model. Our study supports the role of PIEZO1 activation in alleviating *in vitro* and *ex vivo* cystogenesis as a proof of concept. The lack of modulation of in vivo cystic phenotype by collecting duct *Piezo1* knockout reflect redundancy in mechanotransductive pathways, compensation by other cystic nephron segments or off-target effects of Yoda1 in vitro and ex vivo.

## Acknowledgment

The authors would like to thank the Mayo Clinic Center for Cell Signaling in Gastroenterology, particularly the Optical microscopy and microfluidics core (Dr. Mark McNiven and Mr. Eugene Krueger) for their expertise and resources in live cell imaging; Dr. Steven Kleene and Dr. Nancy Kleene for their generous gift of PKD2^-/-^ mIMCD3 cells; and Dr. Timothy Kline for sharing his automated cystic index software.

## Supplemental materials

**Supplemental Fig 1.**
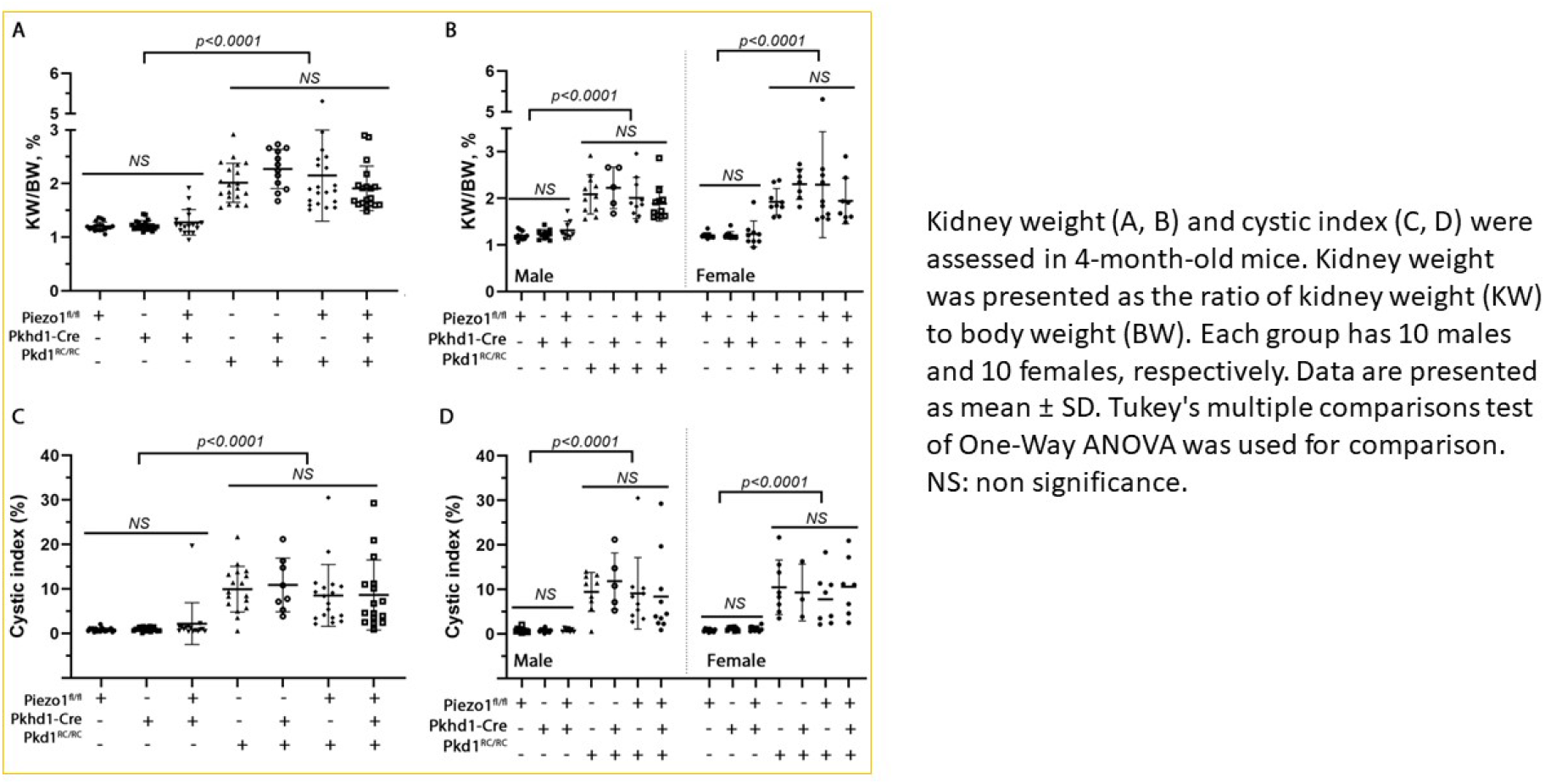
Effects of Piezo1 knockout on kidney weight and cystic index in Pkd1^RC/RC^ mice

**Supplemental Fig 2.**
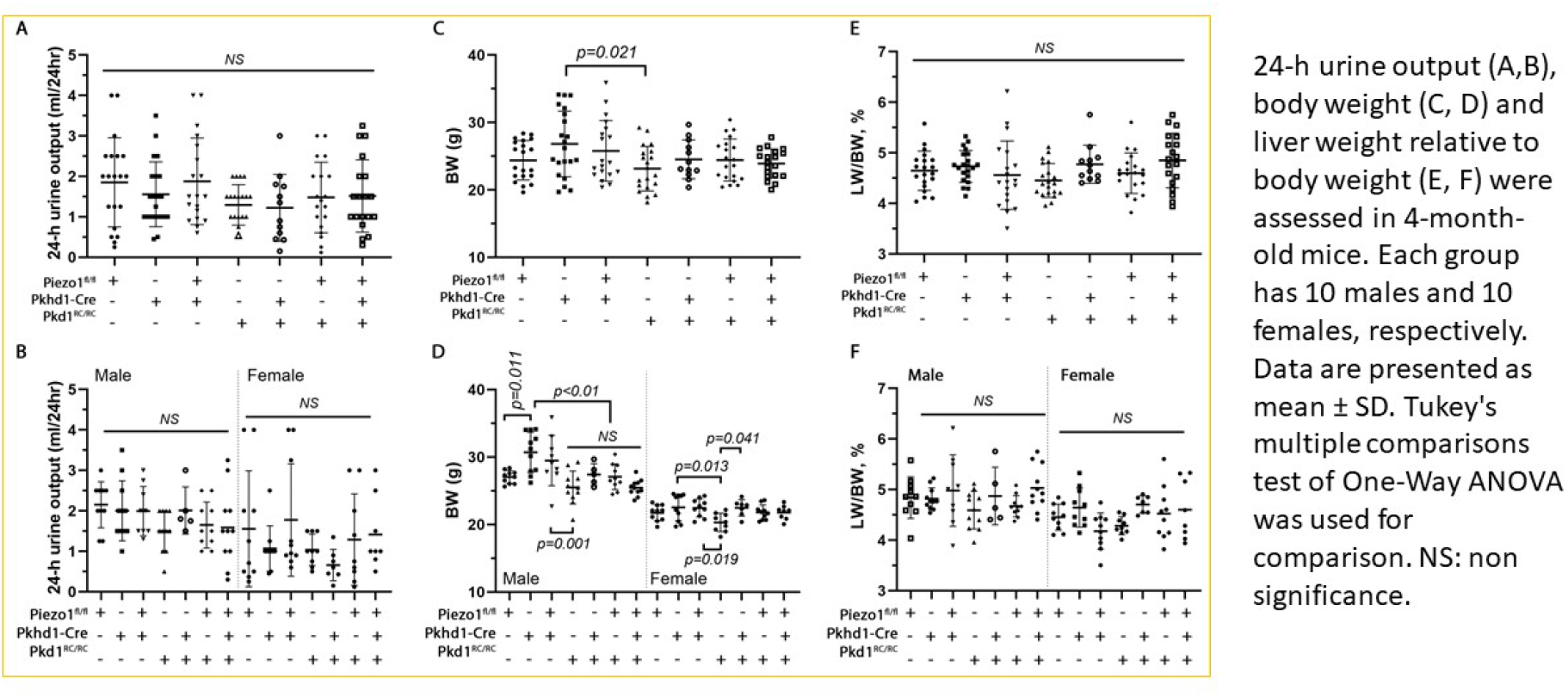
Effects of Piezo1 knockout on 24-hour urine output, body weight and liver weight in Pkd1^RC/RC^ mice

**Supplemental Fig 3.**
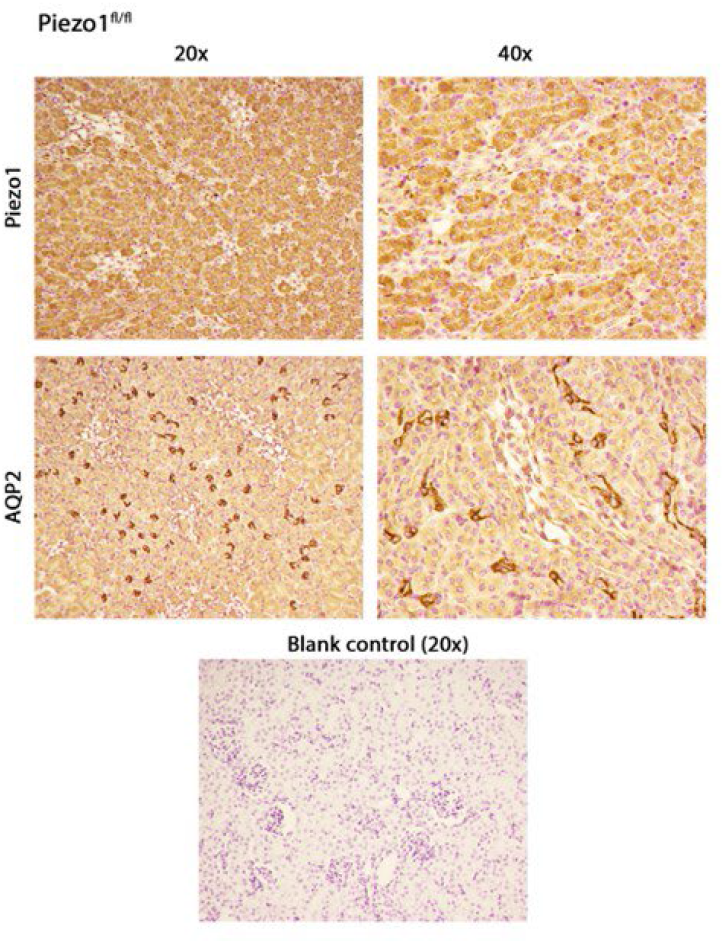
Immunohistochemical staining of PIEZO1 and AQP2 in kidney sections from Piezo1^fl/fl^ mice

**Supplemental Fig 4.**
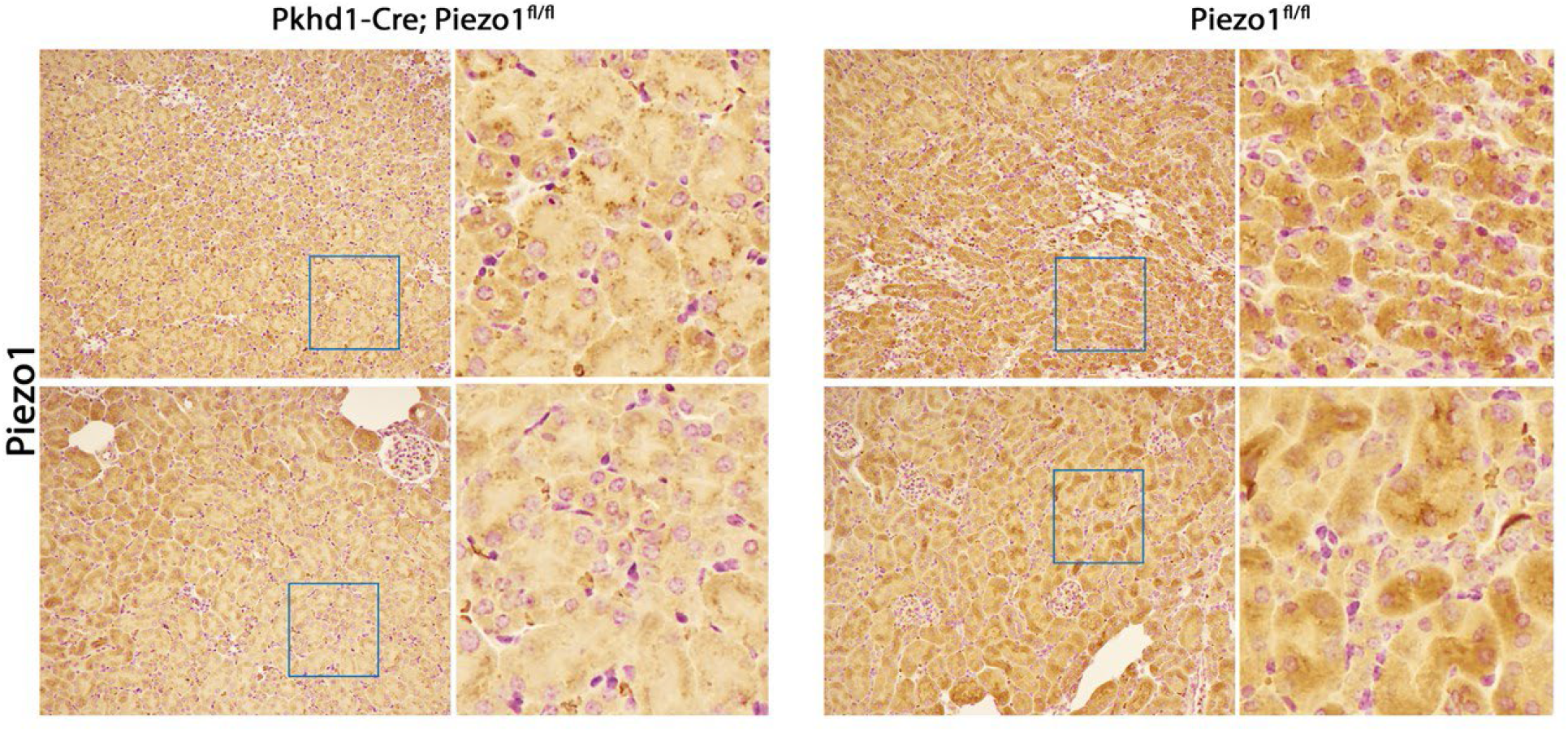
Immunohistochemical staining of PIEZO1 in kidney sections from Piezo1^fl/fl^ and Piezo1^fl/^fl;Pkhd1-Cre mice

**Supplemental Fig 5.**
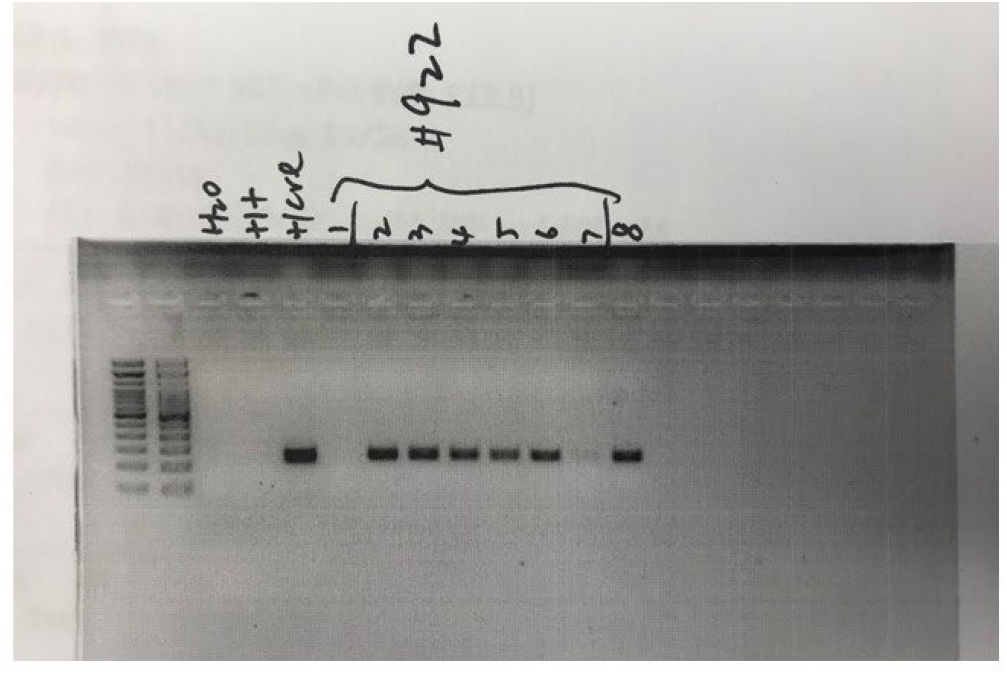
Agarose gel of Cre PCR

**Supplemental Fig 6.**
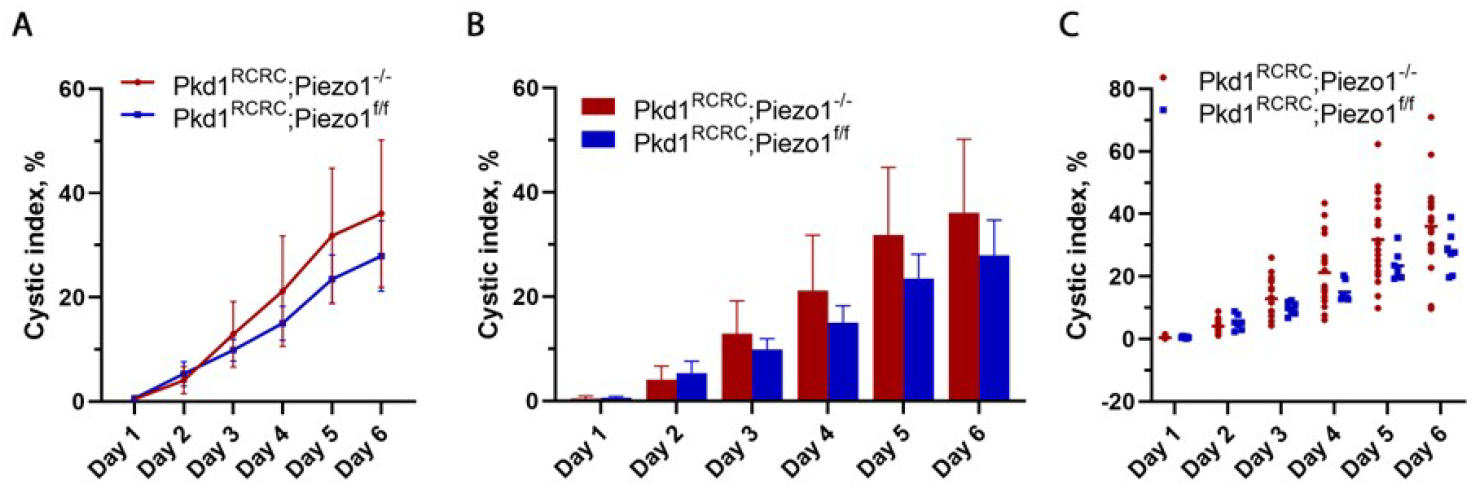
Forskolin-induced *ex vivo* cystogenesis in Pkd1^RC/RC^;Piezo1^-/-^ and Pkd1^RC/RC^;Piezo1^fl/fl^ metanephric organ culture.

